# Period-Specific Remodeling of Luminal Lineages and the Mammary Microenvironment in Dairy Goats

**DOI:** 10.64898/2026.11.20.768023

**Authors:** Mengke Ni, Zhicheng He, Xueqing Han, Yifan Zhu, Xinglong Gong, Wanting Wang, Ping Wang, Jun Luo, Wei Wang, Guoqiang Yi, Cong Li

## Abstract

Milk production requires coordinated remodeling of mammary epithelial lineages and their stromal and immune niches, but this process is incompletely resolved in dairy goats. We aim to define period specific cell states and epithelial niche interactions across the dry to lactation transition. We profile mammary tissue from three peak-lactation and three dry-period Saanen goats using 10× Genomics single-cell RNA sequencing and map one peak-lactation goat with Stereo-seq. Among 44,120 cells, we identify15 populations. Peak lactation is characterized by expansion of luminal alveolar cells and increased β-casein and α-lactalbumin expression, whereas luminal progenitors, hormone-sensing cells, fibroblasts, and B cells are enriched in the dry period. Trajectory inference place luminal progenitors near branches toward alveolar and hormone-sensing states, with sustained *ELF5*-associated activity along the alveolar branch. Myoepithelial, macrophage, natural killer T, and T cell states are also remodeled. SPP1 and MIF signaling networks become more connected and change their cellular targets. These data establish a period resolved atlas of dairy goat mammary remodeling and identify coordinated epithelial, stromal, immune, and communication programs associated with active lactation.

## Introduction

The mammary gland is a dynamic exocrine organ that undergoes repeated cycles of epithelial expansion, differentiation, milk secretion, and postweaning regression (Hannan et al. 2023). In dairy animals, the transition from the postweaning dry period to peak lactation requires coordinated changes in epithelial composition, tissue architecture, and the surrounding microenvironment. These changes are coordinated by endocrine cues and by interactions among luminal cells, myoepithelial (Myoep) cells, fibroblasts (Fibro), vascular cells, and immune populations (Corral et al. 2025). Defining how these compartments are organized across physiological periods is important for understanding lactation biology and improving milk production in dairy animals.

Bulk transcriptomic studies in dairy goats have revealed extensive changes in metabolism, epithelial turnover, immune activity, and tissue remodeling across lactation and the nonlactating period (Xuan et al. 2022). However, whole-tissue profiling averages signals from multiple cell populations and cannot distinguish changes in cellular composition from transcriptional remodeling within individual cell types. At single-cell resolution, the luminal compartment comprises luminal hormone-sensing (HS), luminal alveolar (AV), and luminal progenitor (LP) cells (Li et al. 2025). Myoep cells provide structural and contractile support, whereas Fibro, endothelial (Endo) cells, and immune cells shape the extracellular matrix, vascular interface, and local signaling environment (Kumar et al. 2023). Together, these findings highlight that milk production is a coordinated multicellular process involving both secretory epithelial cells and their surrounding cellular microenvironment (Corral et al. 2025; Yan et al. 2026).

Single-cell and spatial atlases of human and mouse mammary tissue have revealed extensive epithelial, stromal, and immune heterogeneity and established the value of resolving mammary cell states in their anatomical context (Kumar et al. 2023; Li et al. 2020). Recent multi-omic analysis of dairy-goat mammary glands further identified stage-dependent luminal differentiation, regulatory programs, and intercellular communication during lactation and involution (Yan et al. 2026). Nevertheless, how changes in cell abundance and within-cell-type transcriptional states are coordinated across epithelial and niche compartments remains incompletely defined in ruminants (Jaswal et al. 2022).

Here, we used single-cell RNA sequencing (scRNA-seq) to profile mammary tissue from six multiparous Saanen goats, including three at peak lactation and three in the postweaning dry period. We complemented this dataset with spatial transcriptomics from one of the three peak-lactation goats. We examined period-associated changes in cellular composition and transcriptional state, luminal lineage organization, and epithelial–niche communication to define how the mammary gland is remodeled between the dry period and peak lactation.

## Results

### Single-cell and spatial profiling reveal cellular and transcriptional remodeling across lactation states

To characterize cellular dynamics in the dairy goat mammary gland during the dry and peak-lactation periods, we performed 10× Genomics scRNA-seq. After quality control and integration, 44,120 cells were retained and resolved into 15 transcriptionally distinct populations (Figures 1A and S1C). Cell identities were assigned from canonical markers and supported by cluster-enriched transcriptional programs (Figures 1B and S1B, Table S1).

**Figure 1.**
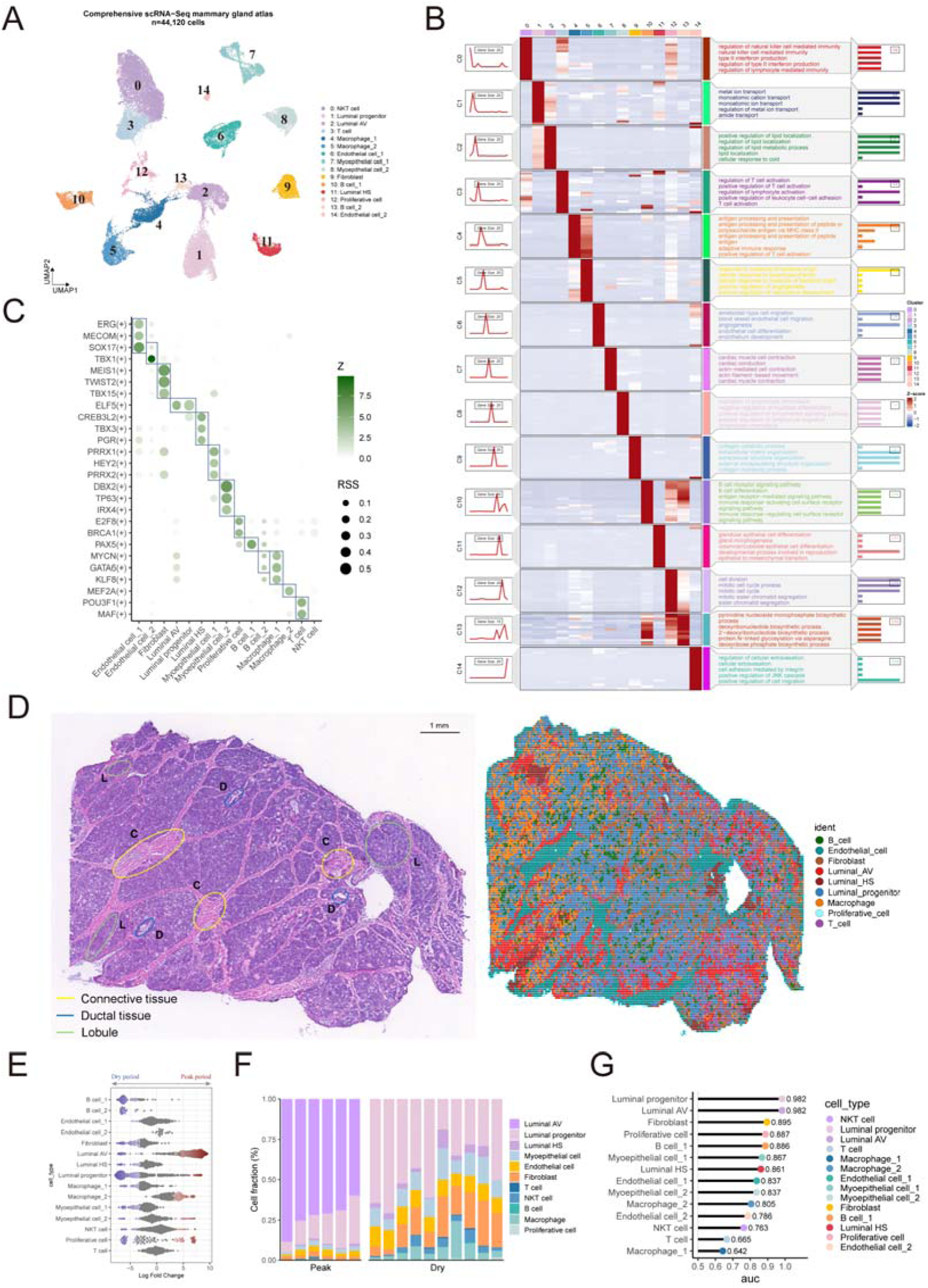
Single-cell and spatial profiling define the cellular and regulatory landscape of the dairy goat mammary gland. (A) UMAP of the integrated scRNA-seq dataset comprising 44,120 cells from six dairy goats, with three biological replicates per period. The 15 populations included LP, AV, HS, Myoep, Fibro, Endo, Mac, B, T, NKT, and Prolif cells. (B) Heatmap of cluster-enriched marker genes and representative Gene Ontology biological processes. (C) Cell-type-specific transcription factor analysis. Transcription factors with τ ≥ 0.8 were retained. Bubble size represents the regulon specificity score, and color represents the corresponding z score. (D) H&E image and spatial cell-type annotation from one section from one peak-lactation animal. Connective tissue (C), ducts (D), and mammary lobules (L) are indicated. Scale bar, 1 mm. (E) MiloR differential abundance analysis of n = 3 biological replicates per period. Each point represents a neighborhood in the k-nearest-neighbor graph; the x axis shows the fold change between peak lactation and the dry period. Neighborhood significance was controlled by the MiloR spatial false discovery rate. (F) Deconvolution-based cell-type fractions estimated from previously generated bulk RNA-seq data (n = 10 dry period and n = 6 peak lactation samples). Cell states were aggregated into their corresponding cell types to construct the single-cell reference signature matrix for deconvolution. Each stacked bar represents one sample. (G) Augur ranking of cell-type transcriptional responsiveness in n = 3 biological replicates per period. AUC indicates the ability to distinguish peak lactation from dry period cells. Spatial results in (D) are descriptive because n = 1 biological replicate. Abbreviations: Myoep, myoepithelial; HS, luminal hormone-sensing; AV, luminal alveolar; LP, luminal progenitors; Prolif, proliferative; Fibro, fibroblasts; Endo, endothelial; Mac, macrophages; NKT, natural killer T cells; T, T cells; B, B cells.

Regulon specificity analysis revealed distinct regulatory programs across mammary cell populations (Figure 1C). Luminal states were separated by ELF5-associated activity in AV/LP and CREB3L2-, TBX3-, and PGR-associated activity in HS, while the two Myoep populations showed distinct PRRX1/PRRX2- and TP63-associated programs. Mac_1 and Mac_2 were likewise distinguished by MYCN/GATA6/KLF8-and MEF2A-associated activity, respectively. Cell-type-specific regulatory patterns were also evident across the remaining stromal and immune populations and were further resolved in the broader regulon-activity heatmap (Figure S1G, Table S2). Together, these profiles provided an independent regulatory layer supporting the cell-type annotations.

We next mapped the inferred cell identities in the single peak-lactation spatial transcriptomic section. H&E staining delineated lobules, ducts, and connective tissue. Luminal populations were concentrated in epithelial-rich regions, Fibro localized to the interstitial compartment, and Endo showed a more distributed, network-like pattern (Figure 1D). Mac, B cells (B), and T cells (T) were more dispersed and locally heterogeneous. Regions selected for dominant epithelial topology still contained variable stromal and immune contributions (Figures S1D-S1F).

Cell neighborhoods differed substantially between the dry and peak lactation period (Figure 1E). AV neighborhoods were enriched at peak lactation, whereas B and Fibro neighborhoods were biased toward the dry period. LP, Myoep, Mac, and Prolif populations contained neighborhoods with varying degrees of period association.

Deconvolution and direct cell count comparisons showed that the two periods differed in global composition and in the relative balance of epithelial, stromal, and immune populations (Figures 1F and S2A-S2D). Augur ranked LP and AV as the most transcriptionally responsive populations, each with an AUC of 0.982 (Figure 1G). Peak lactation was dominated by AV, whereas the dry period contained larger contributions from LP, Fibro, and B. Within the epithelial compartment, AV expanded markedly at peak lactation, whereas LP decreased. Fibro accounted for a larger stromal fraction in the dry period. In the immune compartment, B was more abundant in the dry period, whereas NKT and Mac contributed larger fractions at peak lactation. Thus, the dry to peak transition involved population redistribution together with cell type specific transcriptional remodeling.

### Luminal epithelial populations occupy distinct functional states and an inferred bifurcating trajectory

Marker expression resolved distinct AV, HS, LP, and prolif states within the luminal compartment (Figure 2A). AV cells showed the strongest association with active lactation, with the highest lactation-signature score and a marked expansion from a minor dry period population to the dominant luminal population at peak lactation period (*P* < 0.001; Figures 2B and S3B). Consistent with this shift, peak lactation period AV preferentially engaged lactose/galactose metabolism and protein export pathways, while β-casein and α-lactalbumin signals were significantly increased in peak-lactation mammary glands (*P* = 0.0001 and *P* = 0.0002, respectively; Figures 2C and 2D). In contrast, HS and LP were relatively enriched during the dry period, with LP retaining a stronger progenitor signature (Figure 2E; Figures S3A and S3C). Prolif showed a higher proliferation-signature score at peak lactation without a significant change in abundance (*P* = 0.16; Figure 2F and Figure S3D), indicating increased proliferative activity rather than population expansion.

**Figure 2.**
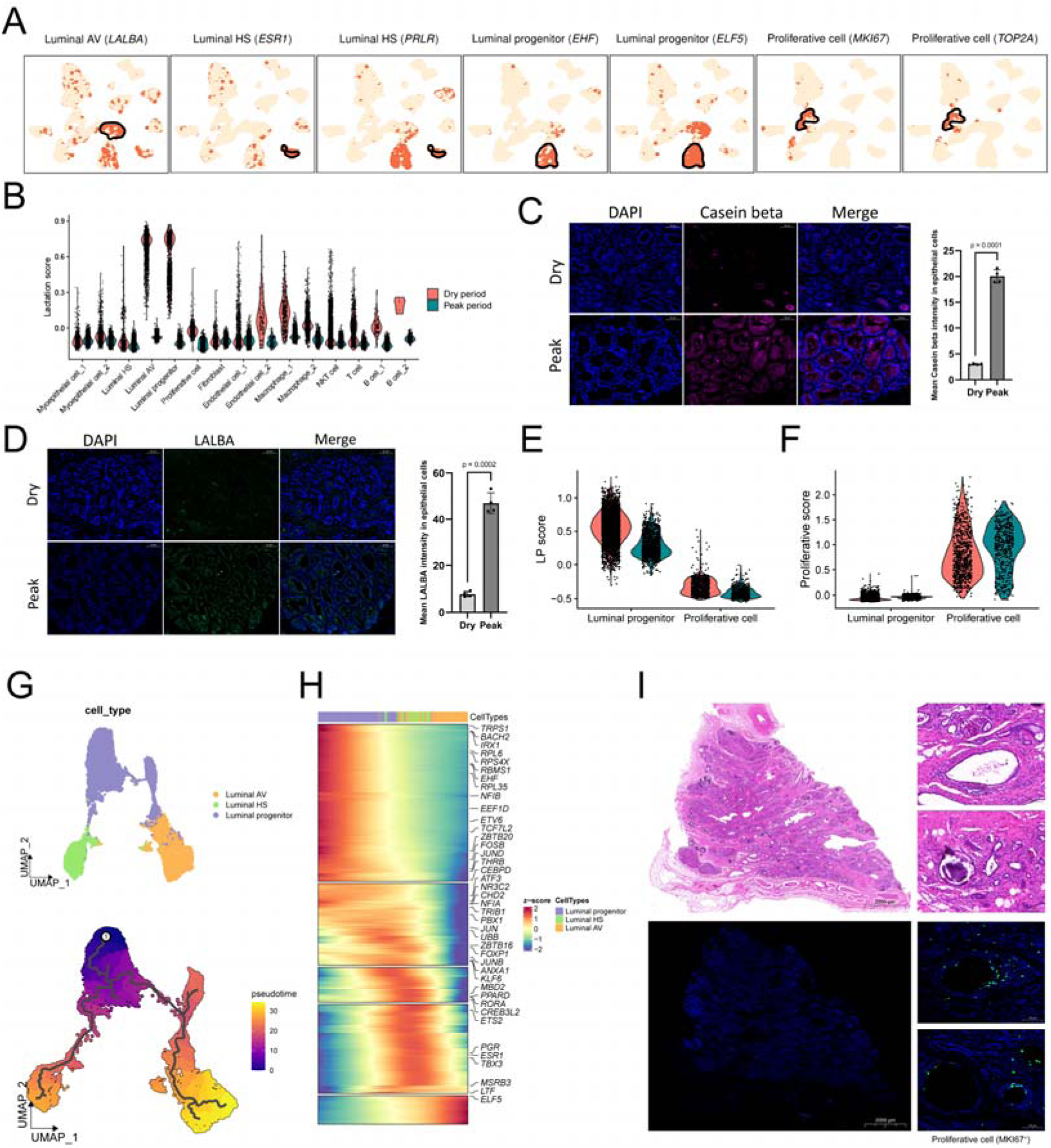
Luminal epithelial populations occupy distinct functional states and an inferred bifurcating trajectory. (A) Feature plots of representative marker genes. (B) Lactation-associated gene-signature scores. (C and D) β-casein and α-lactalbumin expression in dry and peak lactation period mammary tissue. Scale bar, 50 µm. (E) LP-signature scores. (F) Proliferation signature scores calculated from canonical cell cycle genes. (G) Inferred scRNA-seq differentiation trajectory of LP, AV, and HS. (H) Heatmap of 369 pseudotime-associated genes, with cells ordered from low to high pseudotime. (I) *MKI67* expression and spatial distribution in the single peak-lactation mammary section. Scale bars, 2,000 µm and 100 µm. Data are represented as mean ± SEM where indicated; *P* values are from two-tailed Student’s t tests.

CytoTRACE2 predicted that LP exhibited high developmental potency, supporting their designation as the root of the pseudotime trajectory, which subsequently bifurcated into two branches leading toward AV and HS states (Figures 2G and S4C). Spatial trajectory analysis supported a corresponding luminal state transition within the single peak-lactation section (Figure S4A). Along the scRNA-seq trajectory, 369 genes varied with pseudotime, including 41 transcription factors (Figure 2H), with ELF5-associated activity persisting along the AV-directed branch. A linear model incorporating batch, pseudotime, period, and their pseudotime-by-period interaction identified 2,615 genes with period-dependent trajectory dynamics. Of these, 143 increased more steeply with pseudotime at peak lactation, whereas 2,472 showed the opposite interaction pattern. Genes with stronger peak-lactation-associated increases were enriched for translation and peptide biosynthesis, while those showing the opposite pattern were associated with epithelial differentiation, ion transport, cytokine responses, and cellular stress (Figures S4D and S4E). Branch-specific expression further distinguished the LP to AV and LP to HS trajectories (Figures S4F and S4G). In the peak lactation spatial section, MKI67 positive bins were concentrated around duct-associated epithelial regions (Figure 2I), indicating that proliferative activity was spatially localized rather than broadly distributed along the inferred luminal trajectory.

### Myoepithelial subpopulations show complementary structural and contractile programs

Alongside the expansion and secretory activation of AV, efficient lactation also requires myoep contraction to propel milk from the alveoli into the ductal system. We therefore examined how the myoep compartment was remodeled across physiological periods. *MYL9* and *TAGLN* delineated two Myoep populations within the epithelial compartment (Figure 3A). Their relative abundance changed in opposite directions: Myoep_1 was more prominent in the dry period, whereas Myoep_2 increased at peak lactation period (Figure S2B).

**Figure 3.**
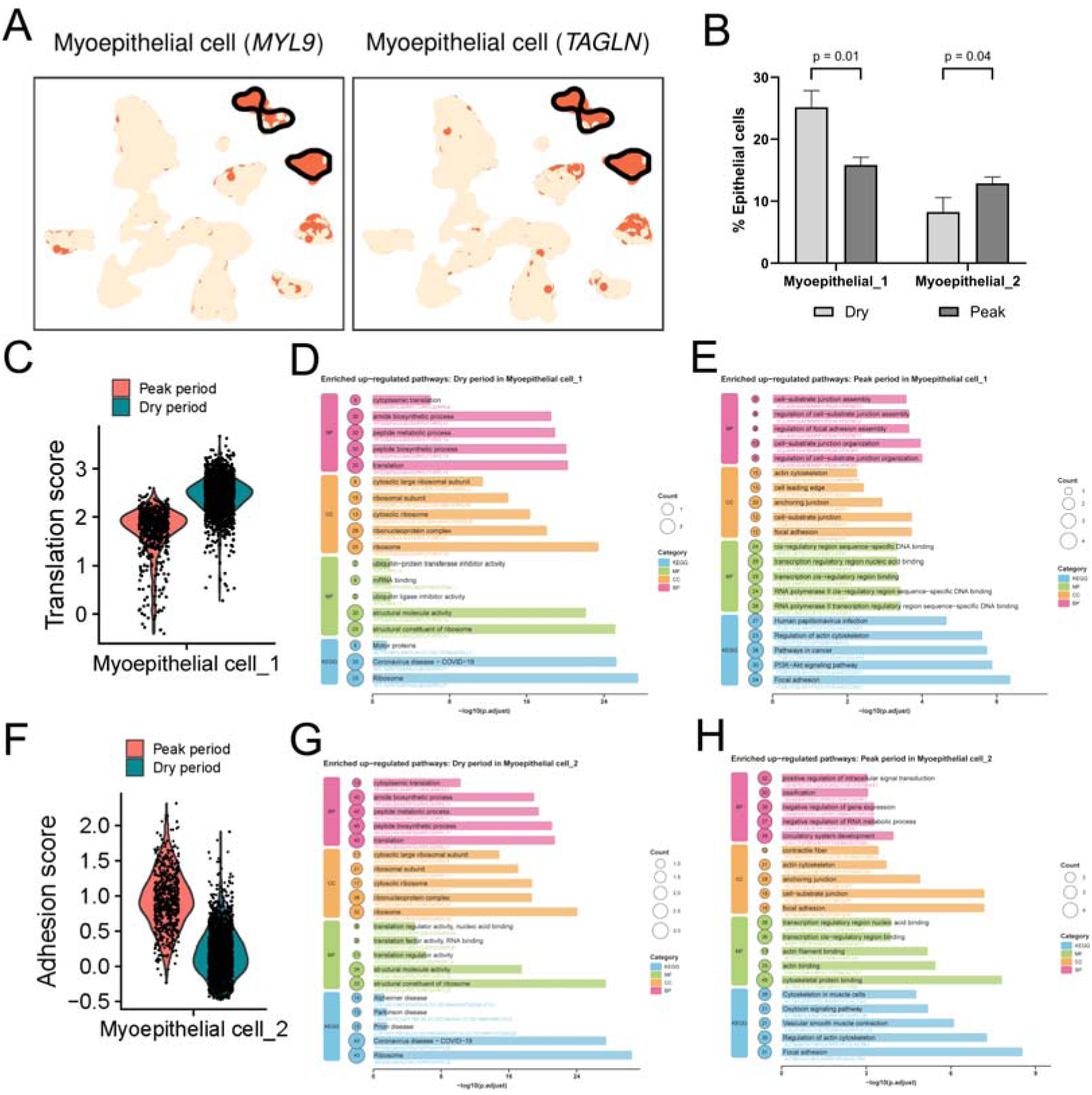
Myoepithelial subpopulations show complementary structural and contractile programs. (A) UMAP feature plots of *MYL9* and *TAGLN* in the epithelial compartment. (B) Relative proportions of Myoep_1 and Myoep_2 during the dry and peak lactation period. (C) Translation module score in Myoep_1. Dots represent individual cells. Functional enrichment of genes upregulated in Myoep_1 during the dry (D) and peak lactation (E) period. (F) Adhesion module score in Myoep_2. Functional enrichment of genes upregulated in Myoep_2 during the dry (G) and peak lactation (H) period, respectively.

Both populations showed translation dominant programs in the dry period. In Myoep_1, dry period genes were enriched for cytoplasmic translation, peptide biosynthesis, ribosomal subunits, and related metabolic processes (Figures 3C and 3D). At peak lactation, Myoep_1 shifted toward cell-substrate junction assembly, focal adhesion, actin cytoskeleton organization, and other adhesion associated programs (Figure 3E). In Myoep_2, the adhesion score was higher at peak lactation than during the dry period (Figure 3F). Dry period Myoep_2 was enriched for translation- and ribosome-related functions (Figure 3G), whereas their peak lactation counterparts were enriched for contractile fibers, actin cytoskeleton, cell-substrate junctions, focal adhesion, and oxytocin-related pathways (Figure 3H). Thus, the two populations shared a biosynthetic state during the dry period but diverged toward adhesion-biased and contractility associated programs during active lactation.

### Stromal compartments undergo distinct period-dependent remodeling

Stromal cell identities were assigned from canonical markers. Fibro expressed *DCN* and *COL3A1*; and Endo cells expressed *PECAM1*. Macrophages (Mac) expressed *TYROBP* and *AIF1*; B cells (B) expressed *CD79A*, *CD79B*, and *MS4A1*. T cells (T) were distinguished by *CD3E* and *CD3G*, and natural killer T (NKT) cells also expressed *NKG7*. Endo_1 and Endo_2 showed little change in relative abundance between periods (*P* = 0.90 and *P* = 0.53, respectively), whereas Fibro was significantly reduced at peak lactation (*P* < 0.001; Figures 4A-4C). Despite the lower Fibro fraction, peak-lactation Fibro was strongly enriched for cell-substrate adhesion, extracellular-matrix organization, ECM-receptor interaction, focal adhesion, and related cytoskeletal programs. Peak-lactation Endo_2 was enriched for collagen-containing ECM, basement membrane, anchoring junction, and focal adhesion. Endo_1 also shifted toward junctional and actin-cytoskeletal programs at peak lactation (Figures S5A-S5D).

**Figure 4.**
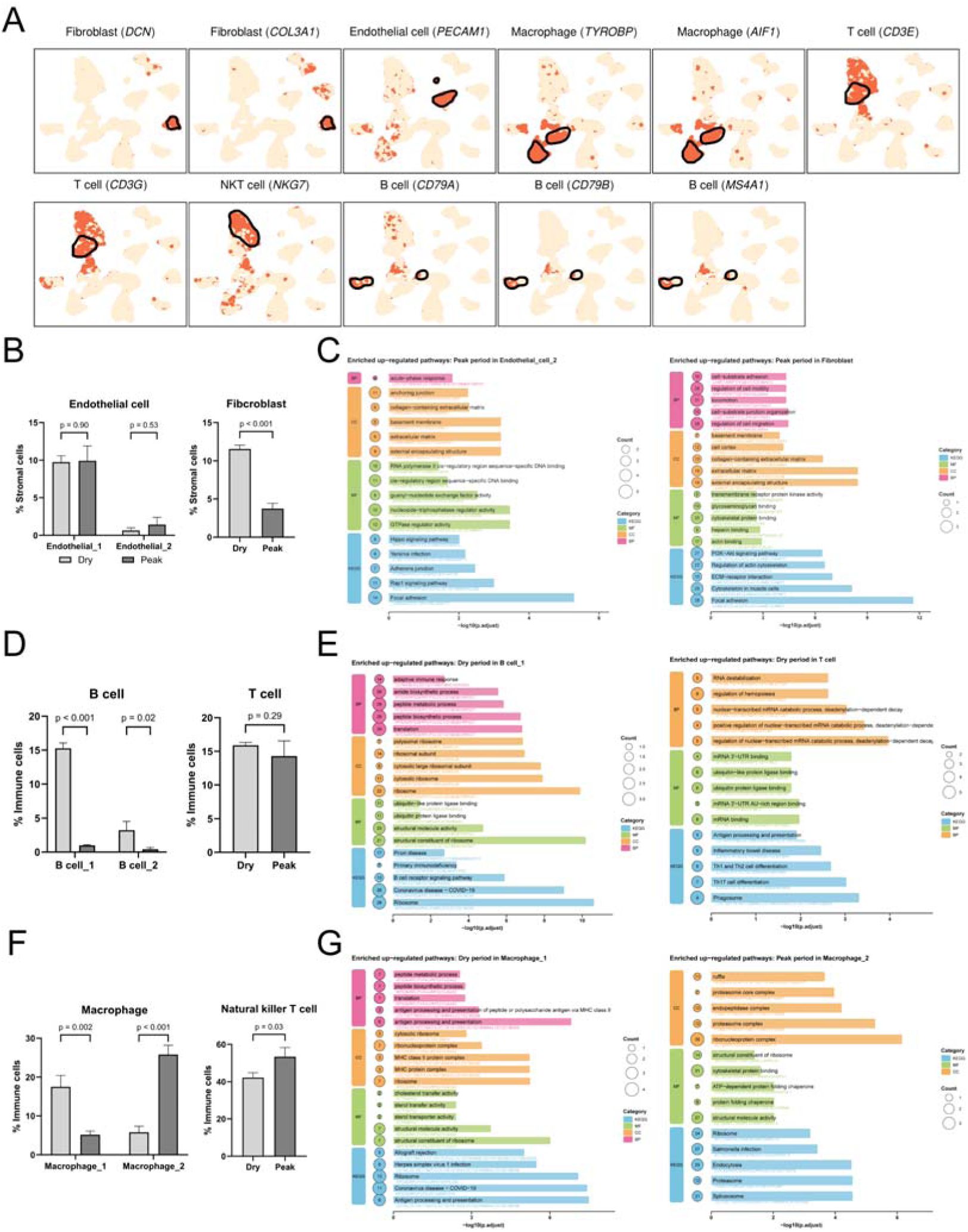
Stromal and immune compartments undergo distinct period-dependent remodeling. (A) UMAP feature plots of representative markers for Fibro (*DCN* and *COL3A1*), Endo (*PECAM1*), Mac (*TYROBP* and *AIF1*), T (*CD3E* and *CD3G*), NKT (*NKG7*), and B (*CD79A*, *CD79B*, and *MS4A1*) cells. (B) Relative proportions of Endo_1, Endo_2, and Fibro within the stromal compartment. (C) Functional enrichment of genes upregulated at peak lactation in Endo_2 and Fibro. (D) Relative proportions of B_1, B_2, and T cells within the immune compartment. (E) Functional enrichment of genes upregulated during the dry period in B_1 and T cells. (F) Relative proportions of Mac_1, Mac_2, and NKT cells within the immune compartment. (G) Functional enrichment of genes upregulated in dry period Mac_1 and peak lactation Mac_2.

B cell abundance decreased at peak lactation, particularly for B_1 (*P* < 0.001), whereas T abundance remained stable (*P* = 0.29; Figures 4D and 4E). Dry period B_1 was enriched for adaptive immune responses and B cell receptor signaling, and dry period T showed antigen processing and presentation together with Th1, Th2, and Th17 differentiation pathways. The residual peak lactation B_2 population showed an acute phase and inflammatory signature (Figures S5E and S5F).

Macrophage populations changed reciprocally. Mac_1 was more abundant in the dry period (*P* = 0.002), whereas Mac_2 increased at peak lactation (*P* < 0.001; Figures 4F and 4G). Dry period Mac_1 showed MHC class II antigen presentation and sterol/cholesterol transport, whereas peak lactation Mac_2 was enriched for endocytosis, proteasome activity, protein-folding chaperones, and cytoskeletal protein binding. Dry period Mac_2 instead showed stronger phagosome and reactive-oxygen species metabolism programs (Figures S5G and S5H). NKT also increased at peak lactation (*P* = 0.03), shifting from antigen-processing and mRNA-regulatory programs in the dry period toward translation associated programs at peak lactation (Figures S5I-S5K).

### Cell communication is rewired around SPP1 and MIF signaling

CellChat-predicted communication networks were more connected at peak lactation than in the dry period (Figures 5A and S6A-S6D). The increase was distributed across epithelial, stromal, and immune populations (Table S3). Myoep_2 gained outgoing communication, Mac_2 showed one of the largest increases in incoming signaling, and Prolif cells changed in both incoming and outgoing activity.

**Figure 5.**
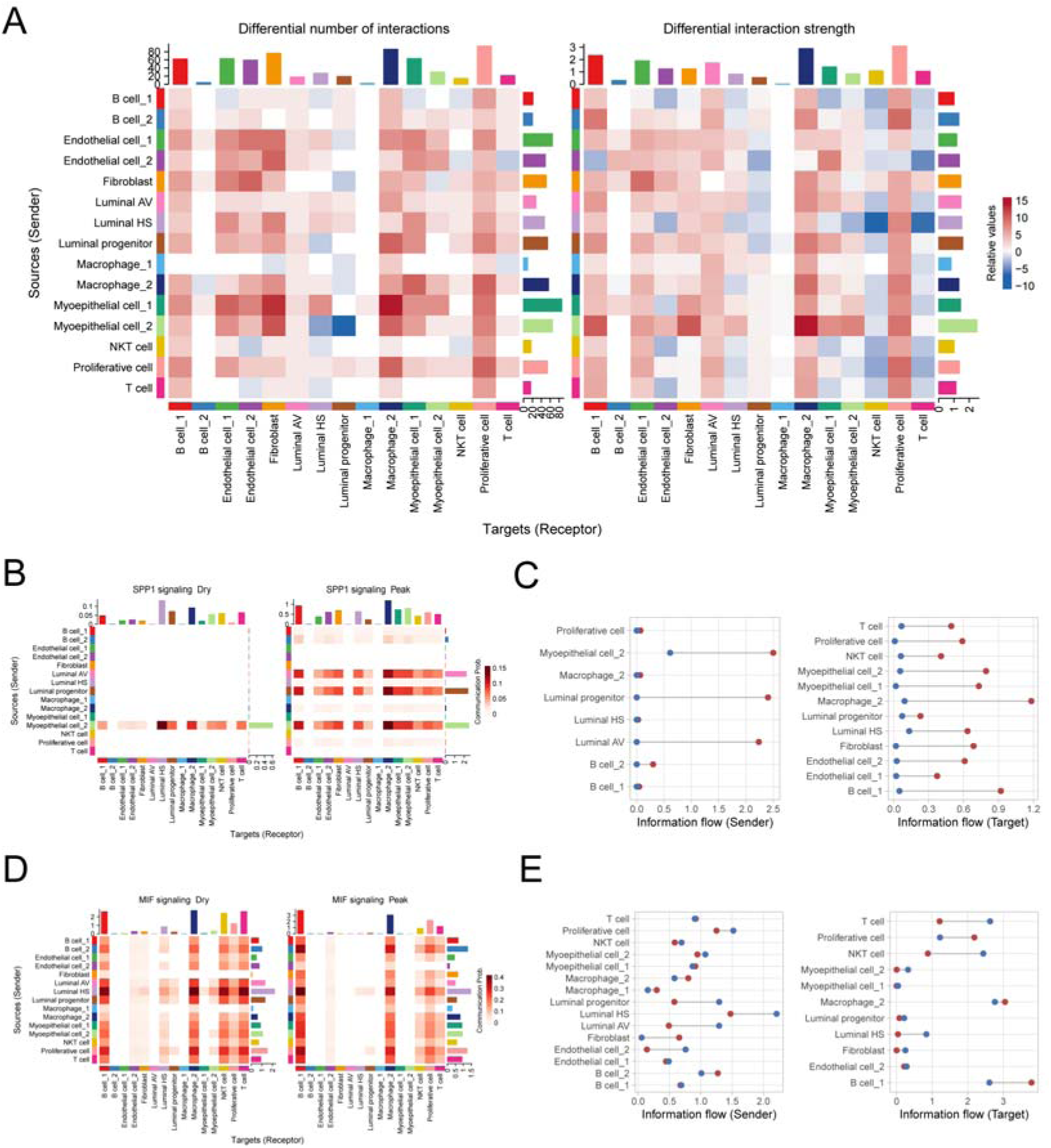
Cell communication networks are reorganized around SPP1 and MIF signaling. (A) Differences in the number and strength of inferred cell-cell interactions between peak lactation and the dry period. Red indicates higher communication at peak lactation period, and blue indicates higher communication during the dry period. Marginal bars summarize changes in incoming and outgoing communication. (B) CellChat-inferred SPP1 signaling network. (C) Outgoing and incoming SPP1 information flow. Red and blue indicate peak-lactation and dry-period cells, respectively. Cell types with zero information flow in both periods were excluded. (D) CellChat-inferred MIF signaling network. (E) Outgoing and incoming MIF information flow. Analyses included n = 3 biological replicates per period. Significant ligand-receptor interactions were identified by a one-sided permutation test with 100 permutations (*P* < 0.05).

Among these changes, SPP1 signaling network showed one of the largest increases in network activity at peak lactation (Figures 5B, 5C, and S6E). In the dry period, SPP1 communication was sparse and largely maintained by Myoep_2. At peak lactation, AV, LP, and Myoep_2 became the main predicted SPP1 sources, with broad connections to epithelial, stromal, and immune populations. Mac_2 showed the strongest increase in inferred SPP1 signaling network as a receiver. B_1, both Myoep, HS, and Prolif also received stronger predicted SPP1 signaling network. These results place secretory luminal cells and Myoep_2 near the center of a peak-lactation SPP1 signaling network.

MIF showed a different pattern of period dependence (Figures 5D, 5E, and S6F). In the dry period, HS, AV, LP, and Endo_2 were prominent predicted sources, and T and NKT received stronger MIF input than at peak lactation. At peak lactation, MIF signaling network remained detectable, but its predicted targets shifted toward B_1, Prolif, and Mac_2. B_1 received stronger MIF input despite its lower abundance. Thus, MIF was not specific to the dry period. Instead, its communication topology shifted from a luminal/Endo to lymphocyte pattern in the dry period to a B, Prolif, and myeloid pattern at peak lactation.

## Discussion

The transition from the dry period to peak lactation was accompanied by coordinated remodeling of the mammary epithelium and its cellular niche. This remodeling centered on the luminal compartment: the dry gland contained a more heterogeneous mixture of LP and HS states, whereas peak lactation was characterized by pronounced AV expansion and increased epithelial expression of β-casein and *LALBA*. These changes indicate that peak lactation entails both redistribution of luminal cell states and acquisition of the biosynthetic program required for milk production. This finding is consistent with single-cell studies showing that luminal epithelial states change in abundance and transcriptional program across mammary physiological periods (Nyquist et al. 2022; Pal et al. 2021). The dry period should therefore be considered a distinct remodeling state in which luminal lineage composition and regenerative programs are reconfigured in ways that may influence subsequent secretory differentiation (Capuco and Choudhary 2020).

Trajectory analysis positioned LP cells upstream of both AV and HS states, supporting a continuous luminal differentiation framework. LP cells remained distinct from Prolif cells, indicating that their persistence does not simply reflect expansion of undifferentiated cycling cells. Instead, they may represent a lineage-biased progenitor state that sustains AV production during established lactation (Nguyen et al. 2018). Mouse single-cell studies have distinguished gestation-associated alveolar progenitors (Avp-G) from lactation-associated alveolar progenitors (Avp-L), demonstrating that the progenitor compartment itself changes across reproductive periods (Bach et al. 2017). On the basis of their physiological context and position along the inferred AV trajectory, the LP cells detected at peak lactation may represent an Avp-L-like state. This interpretation remains provisional because transcriptional similarity across species does not establish equivalent lineage potential.

The expression and inferred activity of *ELF5* and *EHF* provide a regulatory basis for the LP-to-AV transition (Bach et al. 2017). Both factors were associated with LP cells, whereas *ELF5* activity was sustained along the trajectory toward AV differentiation. Previous studies similarly identified *ELF5* and *EHF* as secretory-lineage-associated transcription factors and showed that *ELF5* expression is maintained and further increased as LP cells acquire an alveolar identity (Nightingale et al. 2024; Zhou et al. 2005; Chakrabarti et al. 2012; Bhat-Nakshatri et al. 2024). Together, these observations are consistent with an ETS-family-associated program in which *EHF* may contribute to secretory competence in LP cells, whereas sustained *ELF5* activity supports alveolar specification and activation of the lactogenic program (Oakes et al. 2008). The strong β-casein and *LALBA* signals at peak lactation are consistent with downstream maturation of this lineage (Twigger et al. 2022; Pervolarakis et al. 2020). The epithelial transition occurred together with remodeling of Myoep, Fibro, Endo, and immune populations (Li et al. 2020; Yan et al. 2026). Mac_1 decreased at peak lactation, whereas Mac_2 became the predominant Mac state. Because these populations also exhibited distinct transcriptional programs, this change is more appropriately interpreted as a redistribution of Mac states than as a uniform increase in Mac activity. The antigen-processing and MHC class II-associated features of dry-period Mac_1 is consistent with an antigen-presentation and surveillance state, whereas the endocytic and proteostatic programs of Mac_2 may support material processing and cellular homeostasis in the metabolically active lactating gland. Mammary Mac heterogeneity is not adequately represented by the conventional M1/M2 framework, and the present data do not establish a direct Mac_1 to Mac_2 transition (Li et al. 2020; Dawson et al. 2020; Cansever et al. 2023). Their reciprocal abundance nevertheless indicates that immune remodeling is integral to the dry to lactation transition.

The SPP1 communication network provides a predicted link between Mac remodeling and epithelial adaptation. SPP1 signaling expanded markedly at peak lactation, with AV, LP, and Myoep_2 cells emerging as prominent predicted sources and Mac_2 as a major recipient. This architecture places SPP1 at the interface of epithelial, Myoep, and immune compartments and is consistent with previous single-cell studies identifying SPP1-centered communication in the lactating mammary gland. Given the established roles of SPP1 in integrin- and CD44-associated cell adhesion, cell survival, and extracellular-matrix remodeling, increased SPP1 communication may coordinate mammary epithelial expansion with adaptation of the surrounding niche (Zhang et al. 2026; Hoeft et al. 2023; Li et al. 2026; Muchlińska et al. 2022). In particular, predicted signaling from AV and LP cells to Mac_2 suggests a mechanism by which the secretory epithelium could modulate Mac states or functions during peak lactation. Because several epithelial populations were also predicted to receive SPP1 signals, the same network may provide feedback that contributes to epithelial organization or state maintenance.

MIF signaling showed a context-dependent tissue-maintenance and stress-response pattern. Rather than increasing uniformly, the network was redirected across cellular targets: MIF reception was more prominent in HS and T/NKT during the dry period but shifted toward Mac_2, Prolif, and B_1 at peak lactation. MIF is abundant in human milk, including the aqueous phase and milk-fat-globule fraction, placing the ligand within the lactational environment.(Magi et al. 2002) Extracellular MIF binds CD74 and can signal through receptor complexes involving CD44 (Leng et al. 2003; Shi et al. 2006). Consistent with this pleiotropy, MIF-CD74/CD44 signaling promotes B cell survival and proliferation, whereas MIF-CD74 signaling can support epithelial regeneration after injury through ERK/AKT activation (David et al. 2022; Farr et al. 2020). The period-dependent MIF network may therefore reflect redistribution of cellular competence to respond to a broadly available stress or trophic cue rather than a simple change in total pathway activity. Increased targeting of Mac_2 and Prolif may contribute to immune adaptation and epithelial turnover, whereas reduced targeting of HS and T/NKT is consistent with reorganization of the homeostatic signaling network as the gland enters full secretory activity. In this framework, SPP1 is more closely aligned with adhesion and matrix organization, whereas MIF may redistribute survival and repair associated signaling across epithelial and immune niches. Their complementary organization supports coordinated epithelial immune communication at peak lactation.

Several limitations should be considered when interpreting our findings. Cell-type proportions estimated by scRNA-seq may be influenced by tissue dissociation and compositional effects. The two physiological endpoints do not resolve the temporal sequence of cell-state transitions, and trajectory inference cannot establish lineage relationships. Second, cross-species comparison with published Avp-L signatures, perturbation of *ELF5* and *EHF*, and organoid or *in vivo* manipulation of SPP1 and MIF signaling will be needed to distinguish regulatory drivers from transcriptional correlates. Spatial transcriptomic sampling was limited to one tissue section from a single peak-lactation animal that was also included in the scRNA-seq cohort. These data provide descriptive spatial context but do not constitute independent biological replication or support group-level inference.

This study shows that the dry to peak-lactation transition in dairy goats is a coordinated reorganization of epithelial lineages and their surrounding stromal and immune niches, rather than a simple increase in secretory activity. The expansion and maturation of alveolar cells, the positioning of luminal progenitors along an *ELF5* associated alveolar trajectory, and the reciprocal remodeling of fibroblast, macrophage, lymphocyte, and myoepithelial states together define a period-specific tissue program. The rewiring of SPP1 and MIF communication further indicates that epithelial adaptation is coupled to niche signaling. By resolving these relationships in a ruminant model, this atlas provides a framework for interpreting mammary cell-state plasticity and for developing strategies to improve lactational efficiency and milk traits.

## Methods

### Animals and tissue collection

Mammary gland tissue was collected from six randomly selected, healthy, multiparous Saanen dairy goats maintained at the Northwest A&F University experimental dairy goat farm. All animals were in their second parity and housed under comparable feeding and management conditions. Three goats were sampled at peak lactation period (60 days postpartum), and three goats were sampled during the dry period (300 days postpartum). Individual animals were the biological replicates (n = 3 per period). All procedures were approved by the Institutional Animal Care and Use Committee of Northwest A&F University, China (approval no. DK2024017). After local anesthesia of the mammary gland with procaine, tissue was collected surgically, and visible blood, adipose tissue, and connective tissue were removed with sterile instruments. Samples for scRNA-seq were placed on ice and processed within 30 min. Samples for histology and immunofluorescence were fixed in 4% paraformaldehyde for 48 h. For spatial transcriptomics, an OCT-embedded sample from one peak-lactation goat already included in the scRNA-seq cohort was stored at −80°C until sectioning.

### Preparation of single-cell suspensions

Fresh mammary tissue was minced into approximately 2 mm^3^ fragments and digested in DMEM/F12 (Procell, PM150312) containing 10% fetal bovine serum, 2 mg/mL collagenase XI (Sigma-Aldrich, C9407), and 0.1 mg/mL hyaluronidase (Sigma-Aldrich, H3506) at 37°C for 90 min with agitation. The suspension was further dissociated with 1 U/mL dispase (STEMCELL Technologies, 07913) and 0.1 mg/mL DNase (STEMCELL Technologies, 07470), then passed through a 40 µm cell strainer. Cells were collected by centrifugation at 300 × g for 5 min. Erythrocytes were lysed with red blood cell lysis buffer (BioLegend, 420301), and dead cells were removed with the Dead Cell Removal Kit (Miltenyi Biotec, 130-090-101). The final suspension was washed and resuspended in PBS containing 0.04% BSA. Cell concentration and viability were measured with a Bio-Rad cell counter (1450102). Samples with at least 85% viability were retained for library preparation. Each animal was processed independently through cell isolation, library preparation, and sequencing.

### Single-cell RNA sequencing library preparation and sequencing

Single-cell suspensions were loaded onto the 10× Genomics Chromium platform using the Chromium Single Cell 3′ Reagent Kit v3. Library quality was assessed with an Agilent Bioanalyzer High Sensitivity chip, and libraries were sequenced on a NovaSeq 6000 platform with 150-bp paired-end reads, generating approximately 100 Gb per sample.

### Processing and quality control of scRNA-seq data

Raw scRNA-seq data were processed with Cell Ranger (v7.0.0) using the Capra hircus ARS1.2 genome as the reference. Gene-expression matrices were analyzed in Seurat (v5.0.1), and ambient RNA contamination was removed with decontX (v1.0.0). Cells with 200 < nFeature_RNA < 7,000 and a mitochondrial transcript fraction (percent.mt) < 10% were retained. Putative doublets were identified and removed with DoubletFinder (v2.0.4). After quality control, the six samples were merged and processed with the standard Seurat workflow. Batch effects were corrected with Harmony (v1.2.0). Clustering used FindClusters at a resolution of 0.2, and UMAP visualization used the first 20 Harmony-corrected dimensions. Cluster-specific markers were identified with FindAllMarkers using a Wilcoxon rank-sum test (only.pos = TRUE, min.pct = 0.25, logfc.threshold = 0.25). Cell clusters were annotated manually from established marker genes reported in previous studies.

### Cellular composition analysis

Cell numbers were quantified for each cell type in each individual animal. Cell-type proportions were calculated separately for each biological replicate, using the corresponding parent cellular compartment as the denominator where appropriate. Differences in cell-type proportions between the dry and peak-lactation periods were assessed using two-tailed Student’s t tests based on animal-level proportions (n = 3 per group). Data are presented as mean ± SEM. Differential abundance analysis was additionally performed using MiloR (v2.3.1) based on a neighborhood graph constructed from the 20 Harmony-corrected dimensions. Default MiloR parameters were used, and neighborhood-level significance was assessed with spatial false discovery rate correction.

### Deconvolution analysis

To evaluate cellular composition with complementary bulk transcriptomic data, deconvolution was performed with DWLS (v1.0). Cell type specific count matrices derived from the scRNA-seq dataset were used to construct a signature matrix, which was subsequently applied to bulk RNA-seq datasets including 10 dry period samples and 6 peak lactation period mammary gland samples. The estimated cell-type proportions were then compared with the cellular composition patterns inferred from scRNA-seq analysis.

### Cell-type prioritization analysis

Augur (v1.0.3) was used with default parameters to prioritize cell types according to their transcriptional responses between periods. The area under the receiver operating characteristic curve (AUC) quantified how accurately period-associated transcriptional changes classified cells of each type. Higher AUC values indicated stronger transcriptional differences between periods.

### Differential expression and functional enrichment analyses

To account for inter-animal variability and avoid treating individual cells as independent biological replicates, pseudobulk differential expression analysis was performed separately for each cell type using DESeq2. Raw gene-expression counts were aggregated across cells of the same cell type within each individual animal, generating one pseudobulk expression profile per animal for each cell type. The resulting pseudobulk count matrices were analyzed using DESeq2 to identify differentially expressed genes between the dry and peak-lactation periods, with individual animals treated as independent biological replicates (n = 3 per group). Genes with an adjusted P value < 0.05 were considered significantly differentially expressed.

Functional enrichment analyses were performed using clusterProfiler (v4.10.1). Gene symbols were converted to Entrez and Ensembl identifiers using the bitr function and the Capra hircus OrgDb annotation from Bioconductor AnnotationHub (AH115088). Gene Ontology (GO) and Kyoto Encyclopedia of Genes and Genomes (KEGG) enrichment analyses were conducted using enrichGO and enrichKEGG, respectively. The KEGG organism parameter was set to "chx", and the AH115088 annotation database was used for GO analysis. All genes tested in the corresponding cell-type-specific pseudobulk differential expression analysis were used as the background gene set. Terms with an adjusted *P* value < 0.05 were considered significantly enriched.

### Gene-signature scoring

Lactation, luminal progenitor (LP), proliferation, translation, and adhesion scores were calculated using predefined gene sets listed in Table S4 (Kumar et al. 2023; Twigger et al. 2022; Tirosh et al. 2016). Scores were calculated for individual cells using the AddModuleScore function in Seurat. For statistical comparisons, cell-level scores were first averaged within each cell type for each individual animal. The resulting animal-level mean scores were compared between the dry and peak-lactation periods using two-tailed Student’s t tests (n = 3 per group). Data are presented as mean ± SEM, and *P* values < 0.05 were considered statistically significant.

### Gene-regulatory network and regulon analysis

Gene regulatory networks were inferred with the Python package pySCENIC (v0.12.1). One-to-one orthologs between Capra hircus and Homo sapiens were identified with the NCBI Ortholog database. GRNBoost2 was applied to the count matrix and the human transcription factor list (https://resources.aertslab.org/cistarget/tf_lists/allTFs_hg38.txt) to infer transcription factor-target interactions and calculate link-importance scores. Regulons were evaluated with the human cisTarget motif database. Motif enrichment was quantified using the area under the recovery curve and normalized enrichment score to identify high-confidence regulons. AUCell quantified regulon activity in each cell from its gene-expression ranking.

To identify cell type-specific transcriptional regulators, regulon specificity scores (RSS) were calculated for each regulon across cell types. RSS values were standardized within each cell type using z-score transformation: 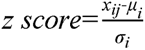; where x_ij_ was the RSS for TF j in cell type i, μ_i_ was the average RSS of all the TFs in cell type i and σ_i_ was the RSS s.d. of all TFs in cell type i. TFs corresponding to regulons ranked among the top three based on z-transformed RSS values and showing high cell type specificity (tau > 0.8) were identified as cell type-specific transcriptional regulators.

### Pseudotime trajectory analysis

The luminal epithelial count matrix was extracted and reprocessed with Seurat for dimensionality reduction, clustering, and visualization. Pseudotime trajectories were inferred with Monocle 3 (v1.3.7) using metadata and dimensionality-reduction coordinates from the Seurat object. CytoTRACE2 (v1.0.0) estimated cellular potency, and the cluster with the highest potency score was selected as the trajectory root.

Genes associated with pseudotime were identified with Monocle 3 graph_test using Moran’s I > 0.25 and q < 0.05. A linear model then evaluated the effects of period, pseudotime, and their interaction, with batch included as a covariate. Genes with significant main or pseudotime-by-period interaction effects (adjusted *P* < 0.05) were selected. The sign of the interaction coefficient was used to classify genes for functional enrichment analysis.

### Cell-cell communication analysis

Cell-cell communication was inferred with CellChat (v2.1.2). Goat gene symbols were mapped to human orthologs, and the secreted-signaling subset of CellChatDB.human was used as the reference. Separate CellChat objects were generated for the dry and peak-lactation periods. Overexpressed genes and ligand-receptor interactions were identified with default parameters, expression was smoothed with PPI.human, and communication probabilities were calculated from normalized data using the triMean method (raw.use = FALSE, scale.distance = 0.01). Significance was assessed with 100 permutations (*P* < 0.05). Pathway-level probabilities and aggregated networks were calculated before comparison of communication number, strength, and pathway activity between periods.

### Stereo-seq library preparation and raw data processing

Spatial transcriptomics was performed on one 10-µm mammary section from a single peak-lactation goat (n = 1 biological replicate); the same animal was one of the three peak-lactation goats profiled by scRNA-seq. An adjacent cryosection was fixed in 4% paraformaldehyde and stained with hematoxylin and eosin for morphological evaluation. The OCT-embedded Stereo-seq section was mounted on a Stereo-seq chip (BGI, China), incubated at 37°C for 3 min, fixed in methanol at −20°C for 40 min, and imaged after nucleic acid staining. After washing, the section was permeabilized with 0.1% pepsin in 0.01 M HCl at 37°C for 5 min. Captured RNA was reverse transcribed on the chip overnight at 42°C. The cDNA was released, purified, amplified, and used to construct a spatially barcoded library, which was sequenced on a DNBSEQ-T7 platform (MGI, China) by Novogene.

FASTQ files were processed with the SAW pipeline. Coordinate identity (CID) sequences were mapped to the designed chip coordinates while allowing one mismatch. Reads were removed when their molecular identifier (MID) contained an N base or more than two bases with quality scores below 10. Retained reads were aligned to the Capra hircus ARS1.2 reference genome, and reads with MAPQ > 10 were assigned to genes. UMIs with the same CID and gene locus were collapsed while allowing one mismatch. The resulting expression matrix was aggregated into non-overlapping Bin 100 units.

### Spatial transcriptomic data processing and annotation

Downstream spatial analyses were performed in R (v4.3.3 and v4.4.1) using Seurat (v5.1.0) and SeuratObject (v5.0.2). The input object contained 21,783 tissue-covered Bin 100 units, with a mean of 1,962 detected genes and 8,821 UMIs per bin, and cluster labels generated at resolutions 1.0 and 1.1. After matching the expression matrix and cluster labels, 21,781 bins were retained. Unless stated otherwise, the Spatial assay was normalized with LogNormalize using a scale factor of 10,000; raw UMI counts were used only for CytoTRACE and Monocle 2 analyses. Cell types were annotated independently at both clustering resolutions by cluster-level gene set variation analysis (GSVA, v1.50.0) using a curated caprine mammary marker panel. A pseudo-bulk matrix was generated for each cluster, and the cell type with the highest GSVA score was assigned to all bins in that cluster. The resolution-1.1 annotations, comprising 13 clusters and nine cell types (AV, HS, LP, Fibro, Endo, Mac, Prolif, T, and B), were used downstream.

The 21,781 annotated bins were divided into a luminal epithelial compartment (LP, AV, and HS; 11,307 bins) and a non-epithelial compartment (10,474 bins). Each subset was reprocessed independently with NormalizeData, FindVariableFeatures (vst method, 2,000 genes), ScaleData, RunPCA (50 components), FindNeighbors, and FindClusters (Louvain algorithm, resolution = 0.6), followed by RunUMAP (30 neighbors, min.dist = 0.3). Marker genes were identified with FindAllMarkers using the Wilcoxon rank-sum test (min.pct = 0.1, logfc.threshold = 0, only.pos = FALSE). Genes with avg_log_2_FC > 0 and Bonferroni-adjusted *P* < 0.05 were considered significant markers.

### Spatial downstream analyses

Cell-cell communication among the nine spatially annotated cell types was inferred with CellChat (v2.1.2) using the same human-ortholog mapping and CellChatDB.human workflow described above. Communication probabilities were calculated from normalized expression values with the triMean method (raw.use = FALSE, scale.distance = 0.01), and significance was assessed with 100 permutations (P < 0.05). Developmental potential was estimated from raw counts with CytoTRACE (v0.3.3) in fast mode for all annotated bins and for the luminal subset. Trajectories were then inferred with Monocle 2 (v2.34.0) using DDRTree. For each analysis, the root was defined as the DDRTree state containing the largest number of bins from the cell type with the highest mean CytoTRACE score. Branch-dependent genes were identified with BEAM, and the 50 genes with the lowest q values at each branch point were visualized in branched expression heatmaps.

Differential expression was tested for all 36 pairwise combinations of the nine annotated cell types with Seurat FindMarkers (Wilcoxon rank-sum test, min.pct = 0.1, logfc.threshold = 0); genes with Bonferroni-adjusted *P* < 0.05 were considered significant. GO and KEGG over-representation analyses were performed with clusterProfiler (v4.5.1) using species-specific Capra hircus gene sets and Benjamini-Hochberg-adjusted *P* < 0.05. Redundant GO terms were grouped with simplifyEnrichment (v1.12.0). Because the spatial dataset comprised one section from one animal, all spatial results were interpreted descriptively and were not used as independent evidence for group-level differences.

### Hematoxylin and eosin staining and Immunofluorescence staining

Mammary gland tissues were fixed in 4% paraformaldehyde for 48 h, dehydrated through a graded ethanol series, and embedded in paraffin. Paraffin sections were deparaffinized, rehydrated, and stained with hematoxylin and eosin (H&E) using standard procedures. After dehydration and mounting, tissue morphology was examined and imaged using Pannoramic MIDI III DX (3DHistech). Paraffin-embedded mammary gland sections were deparaffinized, rehydrated, and subjected to antigen retrieval. After blocking, sections were incubated with primary antibodies against β-casein (MA1-46056, Invitrogen, 1:100), LALBA (YA5359, MCE, 1:100), and Ki-67 (ab16667, Abcam, 1:250), followed by appropriate fluorescently labeled secondary antibodies. Nuclei were counterstained with DAPI, and sections were mounted using an antifade mounting medium. Fluorescence images were acquired using Olympus IX71 under identical imaging settings. Mean fluorescence intensity (MFI) was quantified using ImageJ software to evaluate β-casein and LALBA expression in mammary tissues from the dry and peak-lactation groups.

### Quantification and statistical analysis

Individual animals were considered the biological units of replication for scRNA-seq analyses, with three independent animals per physiological period (n = 3 per group). Cell-type proportions and gene-signature scores were summarized at the animal level before statistical comparisons. Two-group comparisons were performed using two-tailed Student’s t tests by R Stats Package, and data are presented as mean ± SEM. Differential expression and functional enrichment analyses were performed using adjusted *P* values to account for multiple testing, with an adjusted *P* value < 0.05 considered statistically significant. Statistical procedures and significance thresholds for MiloR, CellChat, and trajectory analyses are described in their respective method sections. The spatial transcriptomic dataset comprised one tissue section from one peak-lactation animal (n = 1 biological replicate). Spatial analyses were therefore interpreted descriptively and were not used as independent evidence for group-level statistical inference. All analyses used published software; versions listed in Methods.

## Abbreviations

scRNA-seq: single-cell RNA sequencing
Myoep: myoepithelial
HS: luminal hormone-sensing
AV: luminal alveolar
LP: luminal progenitors
Prolif: proliferative
Fibro: fibroblasts
Endo: endothelial
Mac: macrophages
NKT: natural killer T cells
T: T cells
B: B cells

## Data Access

The data generated in this study have been submitted to the Genome Sequence Archive (GSA) (https://ngdc.cncb.ac.cn/gsa/) under accession number CRA052128. Any additional information required to reanalyze the data reported in this paper is available from the correspondence upon reasonable request.

## Acknowledgements

This work was supported by the National Natural Science Foundation of China (32472874), the General Project of Agricultural Technology Research in Xi’an City (24NYGG0025), and the Key Research and Development Program of Shaanxi Province (2025CY-JJQ-207). The authors declare that the research was conducted in the absence of any commercial or financial relationships that could be construed as a potential conflict of interest.

## Author Contributions

Mengke Ni: Writing - original draft and conceptualization. Zhicheng He: Methodology and software. Xueqing Han: Validation. Yifan Zhu: Formal analysis. Xinglong Gong: Data curation. Wanting Wang: Resources. Ping Wang: Investigation. Jun Luo: Project administration. Wei Wang: Resources and project administration. Guoqiang Yi: Writing - review and editing, supervision. Cong Li: Writing - review and editing, supervision, and funding acquisition. All authors reviewed and approved the manuscript.

## Supplemental information

Document S1. Supplemental Figures S1–S6

Document S2. Supplemental Table S1–S4

Table S1. Cell cluster annotations and canonical marker genes in the dairy goat mammary gland. Table S2. Transcription factor regulon activity across mammary cell populations during the dry and peak lactation period. Table S3. Ligand receptor interactions and signaling pathway information used for cell–cell communication analysis. Table S4. Gene signatures used for functional module scoring in mammary cell populations.

